# The transcriptome and secreted factors of the intervertebral discs in STZ-HFD Type□2 diabetic male mice reveal extensive inflammation

**DOI:** 10.1101/2024.07.31.605332

**Authors:** Christian E. Gonzalez, Rachana S. Vaidya, Sade W. Clayton, Simon Y. Tang

## Abstract

The chronic inflammation observed during type 2 diabetes (T2D) is associated with spinal pathologies, including intervertebral disc (IVD) degeneration and chronic spine pain. Despite the presence of confounding factors, such as obesity, studies have shown that after adjusting for age, body mass index, and genetics (e.g. twins), patients with T2D suffer from disproportionately more IVD degeneration and/or back pain. We hypothesize that chronic T2D fosters a proinflammatory microenvironment within the IVD that promotes degeneration and disrupts IVD homeostasis. To test this hypothesis, we evaluated two commonly used mouse models of T2D – the leptin-receptor deficient mouse (db/db) and the chronic high-fat diet in mice with impaired beta-cell function (STZ-HFD). Compared to their genetic controls—C57BL/6 wild-type mice for STZ-HFD and heterozygous littermates for db/db—STZ-HFD IVDs exhibited more severe degeneration and elevated chemokine expression profiles. RNA-seq further revealed extensive transcriptional dysregulation in STZ-HFD IVDs that was not observed in the db/db model. The STZ-HFD IVDs also expressed enzymes that enhanced production of glycolytic AGE precursors, impaired non-AGE DAMP pathways, and reduced suppressors of RAGE turnover. These results suggest that, under controlled genetic and environmental conditions, the STZ-HFD model more accurately reflect the multifactorial inflammatory milieu characteristic of T2D-induced IVD degeneration.

## Introduction

Type 2 diabetes (T2D) is a prevalent metabolic disorder marked by insulin resistance and prolonged hyperglycemia, impacting millions around the globe and leading to significant healthcare expenses (Srinivasan and Ramarao, 2007; Boucher et al., 2014; Petersen and Shulman, 2018; United States, Center for Disease Control, 2022). This disease shares several characteristics with autoimmune disorders, including the chronic, systemic overexpression of immunomodulating cytokines, which can gradually lead to widespread accrual of tissue damage across multiple organ systems (Itariu and Stulnig, 2014; Chen et al., 2017; de Candia et al., 2019; Daryabor et al., 2020). Among these complications, IVD degeneration is a comorbidity of particular interest due to its chronic-inflammatory etiology (Risbud and Shapiro, 2014; Molinos et al., 2015a; Navone et al., 2017; Lyu et al., 2021; Pinto et al., 2023).

A growing body of research has shown that chronic hyperglycemia, a hallmark of diabetes, initiates and exacerbates degenerative changes in the IVD through several overlapping mechanisms. These include increased apoptosis (Jiang et al., 2013; Kong et al., 2022), pyroptosis (Yu et al., 2023), and autophagy (Jiang et al., 2013) of nucleus pulposus cells, along with matrix degradation mediated by catabolic enzymes such as MMPs and ADAMTSs (Illien-Junger et al., 2013; Kong et al., 2022). Diabetic conditions also suppress anabolic signaling pathways by diminishing expression of growth factors like TGF-β and IGF-1 (An et al., 2017). Furthermore, hyperglycemia-induced oxidative stress and advanced glycation end-product accumulation impair ECM structure and mechanical resilience (Fields et al., 2015; Rosenberg et al., 2023), while reduced VEGFA levels under high-glucose conditions point to disrupted vascular support and nutrient exchange (Wu et al., 2024). Despite these mechanistic connections, the population-level data remains mixed. Fabiane et al., 2016, reported higher LDD scores in individuals with T2D, though significance diminished after adjusting for BMI, and co-twin comparisons were underpowered. Still, biological plausibility remains strong, and Liu et al., 2018, showed that poorly controlled or long-standing T2D correlates with more severe disc degeneration, even after controlling for BMI.

Taken together, these findings suggest that diabetes accelerates IVD degeneration through a multifaceted cascade of molecular and biomechanical alterations. In parallel, epidemiological studies report a strong association between chronic T2D and low back pain (Alpantaki et al., 2019; Cannata et al., 2019; Dario et al., 2017; Jhawar et al., 2006; Liu et al., 2018; Robinson et al., 1998; Sakellaridis, 2006), further implicating T2D as a contributor to spine-related disorders. Chronic inflammation in T2D, driven by a persistent milieu of chemokines, may foster a pro-degenerative microenvironment within the IVD, potentially linking T2D-induced inflammation to accelerated disc degeneration. While this connection is supported by both clinical and experimental data, the precise mechanisms remain poorly understood and warrant further investigation.

Animal models are essential for closely investigating T2D’s impact on IVD degeneration. The db/db mouse model, which harbors a point mutation in the gene encoding for the leptin receptor (Chen et al., 1996; Lee et al., 1996) is widely used to study T2D-related metabolic dysfunctions (Wang et al., 2014). Despite exhibiting many human T2D-like traits—including severe obesity, hyperglycemia, and insulin resistance—its leptin receptor deficiency differs from the multifactorial etiology of human T2D. In spine research, db/db mice show signs of IVD degeneration, such as increased cell apoptosis and extracellular matrix degradation, as evidenced by elevated MMP3 expression and apoptotic markers in the IVD (Li et al., 2020).

Biomechanically, discs from db/db mice demonstrate significantly reduced torsional stiffness and torsion-to-failure strength, indicating potentially compromised mechanical integrity (Natelson et al., 2020). Additionally, db/db discs exhibit histological changes such as increased glycosaminoglycan and collagen content, disrupted NP-AF boundary integrity, and a disorganized collagen fiber network in the annulus fibrosus, despite having comparable AGE levels to wild-type controls (Lintz et al., 2022). However, the lack of leptin signaling could confound interpretations of disc degeneration, since leptin appears to promote anabolic processes and reduce catabolic activities in IVD cells (Curic, 2021; Francisco et al., 2018; Gruber et al., 2007; Han et al., 2018; Li et al., 2013; Segar et al., 2019; Sharma, 2018).

In contrast, the Streptozotocin-High Fat Diet (STZ-HFD) model offers a non-genic approach to replicating T2D. It induces the condition through pro-glycemic diet and low-dose streptozotocin-induced pancreatic beta-cell dysfunction, avoiding the genetic ablation of systemically impactful hormonal pathways like leptin (**Fig. 1A**). This model is characterized by significant metabolic disturbances to glycemic status, insulin resistance, and body weight, mirroring the human T2D phenotype more closely (Kusakabe et al., 2009; Islam and Wilson, 2012). Additional metabolic characteristics reported in STZ-HFD mouse studies include elevated serum insulin levels, dyslipidemia (increased triglycerides, LDL cholesterol, and total cholesterol), and increased markers of inflammation and oxidative stress (Gilbert et al., 2011; Alquier and Poitout, 2018; Yin et al., 2020). This sets this mouse model apart as valuable tool for studying the complex interactions within diabetic complications without the confounding factor of complete leptin signaling ablation (Kusakabe et al., 2009). Previously this model has been employed in studying diabetic complications in bone (Eckhardt et al., 2020). Our study aims to uncover the mechanisms behind inflammatory-pathway contribution to IVD degeneration and dysfunction, advancing the field’s understanding of T2D-related IVD complications.

**Figure 1.**
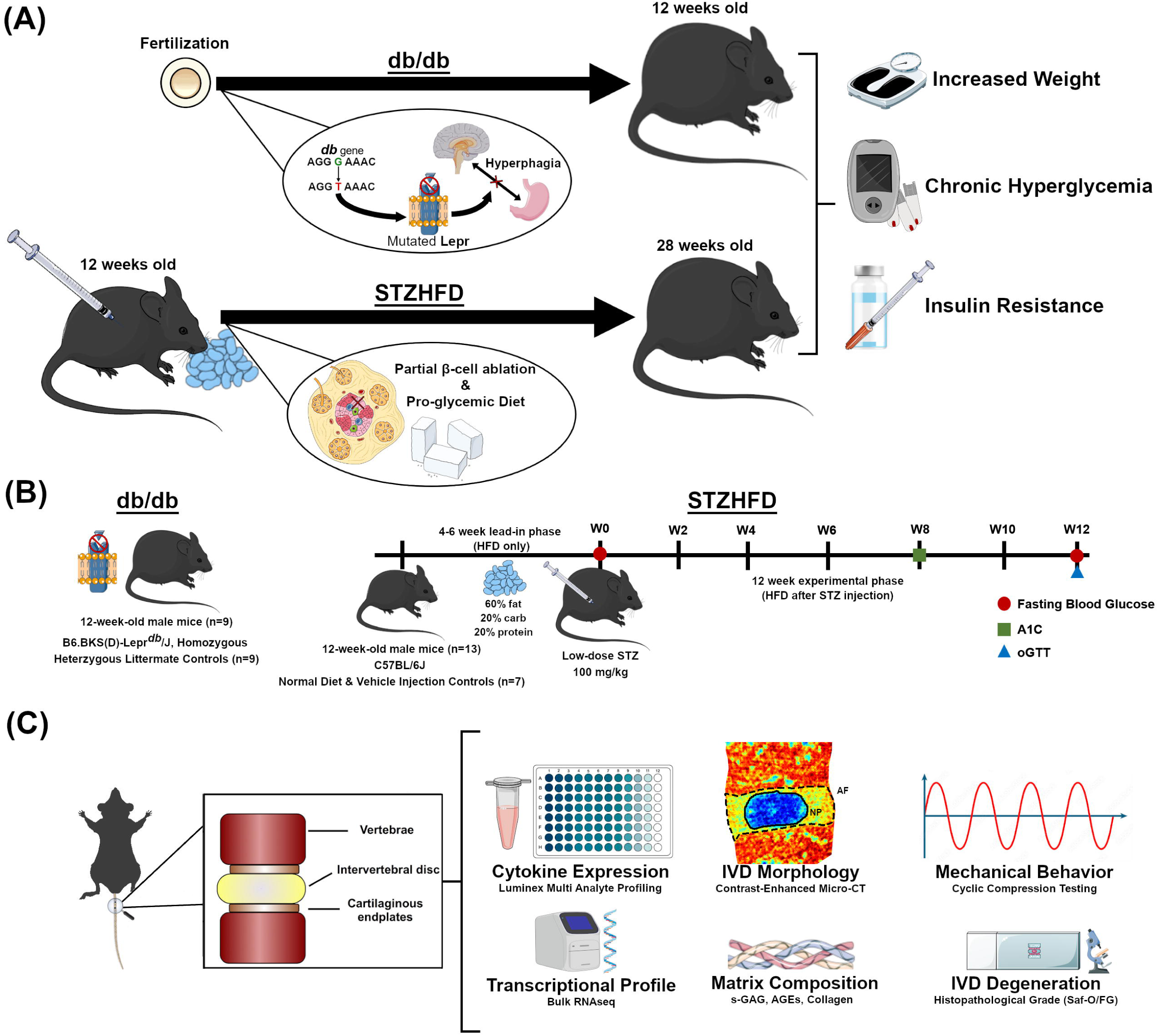
Experimental design, animal model, and workflow of the current study. **(A)** The db/db model arises due to a point mutation in the leptin receptor gene, while the STZ-HFD model develops diabetes through a pro-glycemic diet and beta cell impairment. Both models present symptoms of obesity, chronic hyperglycemia, and insulin resistance, though the magnitude can vary both between and within each model. **(B)** The experimental timeline outlines the progression of the study for both models. The db/db mice are acquired at skeletal maturity (12 weeks old) and sacrificed after metabolic measurements are collected. For the STZ-HFD model, mice undergo a lead-in phase of 4-6 weeks on a HFD followed by a single low dose of STZ. Subsequently, the experimental phase for the STZ-HFD mice continues with HFD for 12 weeks, with periodic assessments of fasting blood glucose, A1c levels, and glucose tolerance. **(C)** After sacrifice, FSUs, consisting of the intervertebral disc and the adjacent vertebral bodies, are extracted from mice. FSUs are then utilized for terminal measures pictured above.

Inflammatory cytokines are pivotal in both IVD degeneration and T2D, mediating acute inflammatory responses while perpetuating chronic inflammation and tissue degradation (Guest et al., 2008; Velikova et al., 2021). They regulate key proteins such as ADAMTs and MMPs, driving the degradation of the IVD’s extracellular matrix (Bond et al., 1998; Malemud, 2019). The identification and characterization of these cytokines in T2D can highlight new therapeutic targets and early intervention markers (Al-Shukaili et al., 2013; Herder et al., 2013). In the spine, inflammatory cytokines contribute to the breakdown of IVD tissue and the development of pain (Risbud and Shapiro, 2014; Shamji et al., 2010). Pro-inflammatory cytokines like TNF-α, IL-1β, and IL-6 are up-regulated in degenerated and herniated disc tissues, exacerbating the inflammatory response and advancing tissue damage (De Geer, 2018; Molinos et al., 2015b; Navone et al., 2017; Wuertz and Haglund, 2013). These cytokines also influence the expression of matrix-degrading enzymes such as ADAMTS-4 and MMP-9, thereby accelerating disc degeneration (Malemud, 2019; Zhang et al., 2015). Their role in T2D is similarly significant, contributing to insulin resistance and beta-cell dysfunction (Calle and Fernandez, 2012). Elevated levels of pro-inflammatory cytokines in patients with T2D underline their importance in disease progression (Guest et al., 2008; Velikova et al., 2021) and targeting these cytokines could open novel therapeutic avenues for both T2D and IVD degeneration (Al-Shukaili et al., 2013; Herder et al., 2013).

This study aims to compare IVD structure and function in db/db and STZ-HFD mouse models of T2D, focusing on inflammation, transcriptomics, and morphological as well as mechanical changes during tissue degeneration (**Figure 1C**) to better understand pathways through which T2D exacerbates IVD degeneration. We hypothesize that the STZ-HFD model, due to its preservation of leptin signaling and induction of systemic metabolic and inflammatory stress, will more accurately capture the chronic inflammatory milieu associated with T2D-driven disc degeneration than the genetically obese db/db model. By evaluating gene expression patterns, inflammatory markers, matrix integrity, and mechanical properties, we seek to delineate the distinct and overlapping pathophysiological features of these two commonly used diabetic models.

This work addresses several unresolved questions in the field: (1) whether the STZ-HFD model better reflects the inflammatory complexity of T2D in the IVD compared to leptin-deficient models; (2) how systemic metabolic stress translates into local disc pathology at molecular and biomechanical levels; and (3) whether chronic hyperglycemia alters DAMP-RAGE interactions and downstream inflammatory signaling in a disc-specific manner. By tackling these gaps, our study contributes a much-needed comparative framework and mechanistic insight into the pathophysiology of diabetic IVD degeneration, while laying the foundation for targeted therapies that address the unique inflammatory profile of T2D-associated disc disease.

## Materials & Methods

### Animals

We used skeletally-mature (12-week-old) male C57BL/6 mice (N = 20) for their established susceptibility to T2D when exposed to a high-fat diet (HFD) and treated with streptozotocin (STZ). Previous findings have indicated that this strain, particularly males, exhibits rapid obesity development under HFD and pronounced insulin resistance following STZ administration (Luo et al., 1998; Mu et al., 2006; Mu et al., 2009). As a result of estrogen-mediated mechanisms of protection, female STZ-HFD mice are resistant to developing T2D and are thus excluded from this study (Medrikova et al., 2012; Pettersson et al., 2012; Stubbins et al., 2012). To contrast the STZ-HFD model’s pathology with a well-characterized model of chronic T2D, parallel cohorts of 3-month-old homozygous (db/db) male Lepr^db^ mutant mice with heterozygous (db/+) littermate controls (n_db/db_ = 9, n_db/+_ = 9) were included, since at this timepoint they have endured a similar duration of T2D symptoms as the STZ-HFD mice. All mice were group-housed (max. 5 mice per cage) under pathogen-free conditions in standard cages; the environment was controlled with a stable temperature and a 12-hour light/dark cycle, with *ad libitum* access to food and water. All procedures were approved by the Institutional Animal Care and Use Committee (IACUC) of Washington University in St. Louis. Regular health and welfare assessments were conducted, including general monitoring of weight, food supply, and behavior.

### Study Design

The study was organized into two phases (**Fig. 1B**). During the lead-in phase (phase 1), mice were maintained on an HFD for four to six weeks, after which they received a one-time dose of STZ. Following injection, mice remained on HFD for an additional 12 weeks during the experimental phase (phase 2). The STZ-HFD group (n = 13) was given a high-fat diet (Research Diets, Inc., D12492i, 60% kcal from fat) for the duration of the study, with the Control + Vehicle group (n = 7) receiving standard mouse chow (5053 PicoLab® Rodent Diet 20, 13% kcal from fat). At the end of the initial phase, baseline measurements of body weight and fasting blood glucose were collected. Following the first phase, STZ-HFD mice were injected intraperitoneally with 100 mg/kg Streptozotocin (MilliporeSigma) in 50 mM sodium citrate buffer (pH 4.5), with Con+Veh animals receiving the sodium citrate buffer only. The two experimental groups received their respective diets for 12 weeks following the injection (experimental phase), and animals were assessed for diabetic status via glucose tolerance. Finally, animals were euthanized, and sterile coccygeal functional spine units (FSUs) including IVDs were harvested from each animal for terminal measurements. For each animal, FSUs were taken from the CC7/8-CC11/12 levels of the tail. Animals were then split between having all IVDs dedicated to RNAseq or to organ culture, mechanical testing, and histology. For animals designated for RNAseq (n_db/+_ = 3, n_db/db_ = 3, n_Con+Veh_ = 4, n_STZHFD_ = 4), the 5 isolated discs were pooled per mouse to isolate sufficient RNA. For animals designated for the other terminal assays (n_db/+_ = 3, n_db/db_ = 3, n_Con+Veh_ = 3, n_STZHFD_ = 9), CC7/8-CC8/9 were used together as pseudo-replicates for organ culture and subsequent histology, CC9/10 was used for contrast-enhanced micro-computed tomography, CC10/11 was used for mechanical testing, and CC11/12 was used for bulk protein assays. In this study we investigate the inflammation and homeostasis in two mouse models of diabetes – the STZ-HFD and the db/db models - by conducting high-resolution molecular profiling and functional phenotyping in the intervertebral disc. Concurrently, the allocation of adjacent FSUs to organ culture, biomechanics, imaging, and protein analysis allows for a comprehensive assessment of structural, mechanical, and biochemical alterations within the same animals, thereby linking molecular changes to disc-level pathology and function in a model-specific manner.

### Measures of diabetic status

Several measures of diabetic status were collected at specific time points during the experimental phase: fasting blood glucose (FBG), glucose tolerance test (GTT), and body weight were measured during week 12, while percentage of glycated hemoglobin (%A1C) was measured at week 8 [**Fig. 1A**]. Blood glucose levels (mg/dL) were measured using a glucometer (GLUCOCARD Vital^®^ Blood Glucose Meter). Blood samples were drawn via superficial incision to the tail tip of fasted mice using a scalpel; the tail was immediately treated with analgesic (Kwik Stop® Styptic Powder) after blood collection. The % A1C was measured using the A1CNow®+ system (PTS Diagnostics) according to kit instructions. Blood samples were drawn fresh in the same way as during the blood glucose test; previously frozen samples from db/db and db/+ animals were excluded due to quality issues, and all analyzed samples included were from fresh blood draws from later cohorts. Finally, for GTT, mice were fasted and had their blood glucose measured as described above to establish a baseline. Mice were then injected intraperitoneally with 2 g/kg glucose in sterile water. Additional blood glucose measurements were taken at 30 min., 60 min., and 90 min. post injection. The area-under-the-curve (AUC) of blood glucose (mg•h/dL) was calculated over the course of the test. For the purpose of evaluating diabetic status for inclusion in the study, a cutoff of 435 mg•h/dL on the GTT, established in previous studies on human T2D criteria (Sakaguchi et al., 2015), was adjusted for time and interspecies differences in blood glucose levels. Discrepancies in sample numbers across measures in the db/db and db/+ groups resulted from poor blood sample quality for A1c measurement or glucometer errors during FBG collection.

### Organ culture

Following extraction, FSUs were cultured in 2 mL Dulbecco’s Modified Eagle Medium/Nutrient Mixture F-12 Ham with L-glutamine and 15 mM HEPES (Sigma-Aldrich, D6421). Culture medium was supplemented with 20% fetal bovine serum (Gibco No. A5256801) and 1% penicillin-streptomycin (Gibco No. 15140122). Cultures underwent a preconditioning period of 7 days with regular media changes to account for the inflammatory response from extraction (Fig. 1C). Conditioned media was collected 48 hours after the final media change at the end of the preconditioning period and immediately frozen in -80° C.

### Chemokine assay

A multiplex assay of remodeling factors and inflammatory chemokines (45-Plex Mouse Cytokine Discovery Assay, Eve Technologies Assays; CCL-2,3,4,5,11,12,17,20,21,22; CSF-1,2,3; IL-1α,1β,2,3,4,5,6,7,9,10,11,12A,12B,13,15,16,17; CXCL1,2,5,9,10; CX3CL1; IFN-γ,β1; TNFα; LIF; VEGF; EPO; TIMP-1) was performed on conditioned media samples. Psuedo-replicates from the same animal were pared down to one representative value by selecting the median cytokine level, and cytokine levels were comparatively analyzed using Welch’s t-Test (n_db/+_ = 3, n_db/db_ = 3, n_Con+Veh_ = 3, n_STZHFD_ = 9). Cytokines with greater than 25% missingness (values outside of assay range) across all experimental groups were excluded from further analysis. Significantly up-regulated cytokines in each model were selected for a secondary fold change analysis, where protein expression levels for db/db and STZ-HFD mice were used to calculate fold change for each cytokine over the corresponding average expression of the control (db/+ and Con+Veh respectively).

### Cytokine Interaction Network Construction and Analysis

To further investigate the inflammatory profiles of these T2D models, networks of cytokine interactions were constructed and analyzed using a custom MATLAB (Version: 9.13.0.2080170 R2022b) script as in previous studies (Easson et al., 2023). Networks were generated by calculating a Pearson correlation matrix for each experimental group based on cytokine expression data from the multiplex panel of conditioned media. To interrogate the strong protein correlations, a threshold (|r| > 0.7) was applied to the correlation matrices. This identifies cytokines that are likely co-regulated, meaning they may be part of the same inflammatory pathway or respond to the same triggers. The filtered matrices were used to create undirected graphs, with nodes representing cytokines and edges representing significant interactions.

Centrality measures were calculated to determine the importance of each cytokine within the networks. Eigenvector centrality and betweenness centrality were computed for each network using the centrality function. Eigenvector centrality aids in identifying cytokines that are dominant in shaping the entire inflammatory response, indicating they are connected to many other active cytokines. Betweenness centrality identifies cytokines that are critical in cross signaling between otherwise separate inflammatory signals. The resulting centrality values were organized into tables and sorted to identify the top-ranking cytokines. For each centrality metric, shared high-ranking cytokines between the diabetic models and unique cytokines for each diabetic model were aggregated. This was used to identify shared and unique inflammatory pathway drivers in each model. Additionally, key network characteristics were extracted to understand the structure and function of the cytokine networks. The average path length was determined using the distances function to compute the shortest finite paths between all pairs of nodes. This can be used to indicate how inflammatory signals spread across the network and the parity among similarly regulated cytokines. Modularity and community structure were assessed using the Louvain community structure and modularity algorithm (Blondel et al., 2008).

Modularity in this context identifies sub-pathways and clusters in the network. The k-hop reachability was computed to assess the extent to which cytokines can influence each other within one (k=1) or two (k=2) degrees of separation. This indicates how far a single cytokine’s influence reaches within the network. The Jaccard index was used to compare the reachability matrices between different groups, providing a direct measure of similarity between networks. Finally, the networks were visualized using force-directed layouts with nodes colored by eigenvector centrality and sized by betweenness centrality.

### Histology

Following removal from culture, FSUs (n_db/+_ = 3, n_db/db_ = 3, n_Con+Veh_ = 3, n_STZHFD_ = 9) were fixed in 10% neutral-buffered formalin (Epredia™ 5735) overnight and decalcified in ImmunoCal (StatLab STL14141) for 72 hours. Samples were embedded in paraffin blocks, sectioned in the sagittal plane at 10 µm thickness, and stained with Safranin-O/Fast Green prior to being imaged via Hamamatsu NanoZoomer with a 20x objective. Blinded histological images of the IVDs were evaluated for degeneration based on a standardized histopathological scoring system (Melgoza et al., 2021).

### Contrast-enhanced Micro-computed Tomography

Functional spine units (n_db/+_ = 3, n_db/db_ = 3, n_Con+Veh_ = 3, n_STZHFD_ = 9) were incubated in a 175 mg/mL Ioversol solution (OptiRay 350; Guerbet, St. Louis) diluted in PBS at 37 °C. Following 4 hours of incubation, the samples underwent scanning with a Viva CT40 (Scanco Medical) at a 10-µm voxel size, using 45 kVp, 177 µA, high resolution, and a 300 ms integration time. CEµCT data was exported as a DICOM file for analysis in a custom MATLAB program (https://github.com/WashUMusculoskeletalCore/Washington-University-Musculoskeletal-Image-Analyses). After an initial Gaussian filter (kernel size = 3), functional spine units were segmented by drawing a contour around the perimeter of the IVD every 10 transverse slices and morphing using linear interpolation. This was defined as the whole disc mask. The NP was segmented from the whole disc by thresholding and performing morphological close and morphological open operations to fill interior holes and smooth NP boundaries. The volumes and intensities were calculated from the NP and whole disc regions. Disc height index (DHI) was measured by averaging the height-to-width ratio of the IVD over five slices in the mid-sagittal plane. Finally, the NP intensity/disc intensity (NI/DI) ratio and NP volume fraction (NPVF; NP volume / total volume) was computed using the intensity and volume metrics reported by the output analysis within the MATLAB program. All thresholding and analysis were performed using blinded and validated methods (Lin et al., 2016; Lin and Tang, 2017).

### Mechanical Testing

Mechanical testing of FSUs (n_db/db_ = 3, n_db/+_ = 3, n_STZHFD_ = 3, n_Con+Veh_ = 9) was performed using cyclic compression on a microindentation system (BioDent; Active Life Scientific) with a 2.39 mm probe as previously described (Liu et al., 2015). Samples were adhered to an aluminum plate and placed in a PBS bath prior to aligning the sample beneath the probe with a 0.03 N preload. Each unit was then sinusoidally loaded in compression at 1 Hz for 20 cycles with a 35 μm amplitude. A loading slope value was calculated from the linear region of the force-displacement curve, and the loss tangent (tan delta) was calculated from the phase delay between loading and displacement (Liu et al., 2015). A representative force displacement curve can be seen in **Supplemental Figure S1**.

### Matrix Protein Assays

Whole extracted discs (n_db/db_ = 3, n_db/+_ = 3, n_STZHFD_ = 3, n_Con+Veh_ = 9) were used to measure the biochemical content of various matrix proteins. First, discs were digested overnight in a papain digestion buffer, after which the buffer was collected for a 1,9-dimethylmethylene blue assay of sulfated glycosaminoglycan content with a chondroitin sulfate standard (Liu et al., 2017). The disc was then subjected to high temperature bulk hydrolyzation in 12 N HCl. Hydrolysates were desiccated and reconstituted with 0.1x phosphate-buffered saline (PBS) and measured against a quinine standard for advanced glycation end product content (Liu et al., 2017). Finally, a hydroxyproline assay was used to quantify collagen content (Liu et al., 2017).

### RNA Collection and Sequencing Analysis

Sterile extracted IVDs (n_db/+_ = 3, n_db/db_ = 3, n_Con+Veh_ = 4, n_STZHFD_ = 4) were placed directly into cold media. After extraction, IVDs were flash-frozen in liquid nitrogen and homogenized via ball mill (Sartorius Mikro-Dismembrator U). Homogenates were resuspended in TRIzol Reagent (Invitrogen 15596-026) and centrifuged at 800 RCF for 5 minutes. Supernatant was collected, purified, and isolated using column filtration (Zymo Research Direct-zol RNA Microprep Kit R2060). Samples were prepared, indexed, pooled, and sequenced on an Illumina NovaSeq X Plus, with basecalls and demultiplexing done using DRAGEN and BCLconvert version 4.2.4. RNA-seq reads were aligned to the Ensembl release 101 assembly with STAR 2.7.9a1, and gene counts were derived using Subread:featureCount version 2.0.32. Isoform expression was quantified with Salmon 1.5.23, while sequencing performance was assessed using RSeQC 4.04. Gene counts were normalized using EdgeR, and low-expressed genes were excluded. The count matrix was transformed to moderated log 2 counts-per-million with Limma’s voomWithQualityWeights, and differential expression analysis was performed. Specific genes related to sugar reduction and dicarbonyl-compound formation (a precursor reaction in the formation of AGEs) were initially identified, along with ligands for RAGE, regulators of NF-kappaB function, and adipokines. A subsequent broader analysis of global perturbations in GO terms, MSigDb, and KEGG pathways was performed using GAGE. To better understand the types of the pathways altered in T2D, pathway analysis was grouped into functional categories for visualization. For shared or similar pathways within each category, the broader pathway encompassing more genes was elected for visualization. All pathways and genes were filtered for significance with corrections for false-discovery rate, and individual genes were excluded if the |log_2_ (fold change)| < 1. All visualization was performed using GraphPad Prism 10.3.1 v509. The data discussed in this publication have been deposited in NCBI’s Gene Expression Omnibus (Edgar, 2002) and are accessible through GEO Series accession number GSE288503 (https://www.ncbi.nlm.nih.gov/geo/query/acc.cgi?acc=GSE288503).

### Statistical Analysis

All statistical analysis was performed in GraphPad Prism 10 software. Comparisons between grouped fold change values (cytokines) were conducted using an Unpaired t-test with Welch’s correction, and comparisons of all groups in other data were conducted using Brown-Forsythe and Welch’s ANOVA with Dunnett’s T3 as post-hoc. Results were considered statistically significant when *p* < .05. P-values were directly reported on each figure where relevant.

## Results

### The db/db and STZ-HFD Mouse Models Exhibit a Characteristically Diabetic Phenotype

Both db/db and STZ-HFD mice demonstrate hallmark features of diabetes (**Fig. 2**). Both groups exhibit AUC GTT values above the defined threshold, indicating impaired glucose tolerance (**Fig. 2A**). The STZ-HFD mice show severe glucose intolerance compared to Con+Veh mice (p < 0.0001). The terminal body weights show that db/db mice weigh significantly more than both db/+ and STZ-HFD mice, (**Fig. 2B**). The terminal fasting blood glucose levels indicate a significant difference only between the STZ-HFD and Con+Veh groups, demonstrating notable fasting hyperglycemia in the STZ-HFD mice, but not in db/db mice (**Fig. 2C**). HbA1c levels reveal no difference between db/db mice and STZ-HFD mice, but each group significantly differs from their respective controls (**Fig. 2D**). This indicates that both models exhibit chronic hyperglycemia, confirming their relevance as models of T2D.

**Figure 2.**
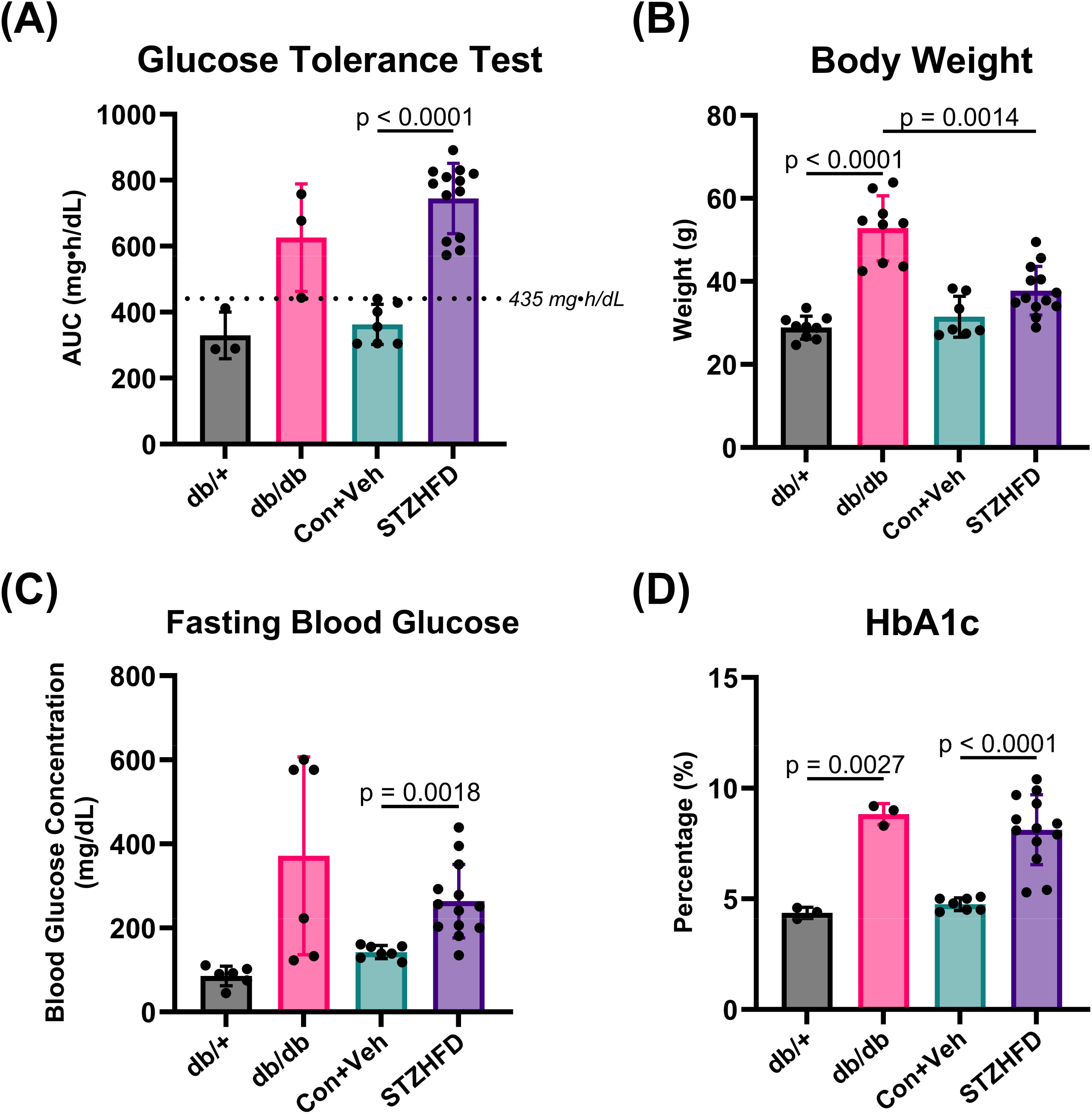
Both db/db and STZ-HFD mice represent a characteristically T2D phenotype. **(A)** AUC GTT shows elevated glucose intolerance in db/db and STZ-HFD mice, with a highly significant difference between Con+Veh and STZ-HFD (p < 0.0001). **(B)** Terminal body weights indicate db/db mice are significantly heavier than db/+ and STZ-HFD mice. **(C)** Terminal fasting blood glucose levels are significantly higher in STZ-HFD mice compared to Con+Veh. **(D)** HbA1c levels indicate chronic hyperglycemia in both db/db and STZ-HFD mice, with no difference between them, but significant differences from their respective controls.

### Histopathological Analysis Reveals IVD Degeneration in STZ-HFD Mice

The IVDs of STZ-HFD mice exhibited more degeneration compared to the IVDs of controls and the db/db mice. **Figure 3** presents a comparison of histological phenotypes across the four groups (db/+, db/db, Con+Veh, and STZ-HFD), showing the range of histopathological scores within this study (**Fig. 3A**). Histopathological scoring revealed that only the STZ-HFD mice exhibit significantly greater IVD degeneration compared to the control group (**Fig. 3B**). In the Con+Veh samples, the annulus fibrosus has healthy, convexed outer lamellae (i+) and well-organized, concentric inner lamellae (T). In contrast, the STZ-HFD samples show degenerate crimped, concave outer lamellae (¢) and wavy, disorganized inner lamellae (V), indicating altered matrix structure in the AF (**Fig. 3C**).

**Figure 3.**
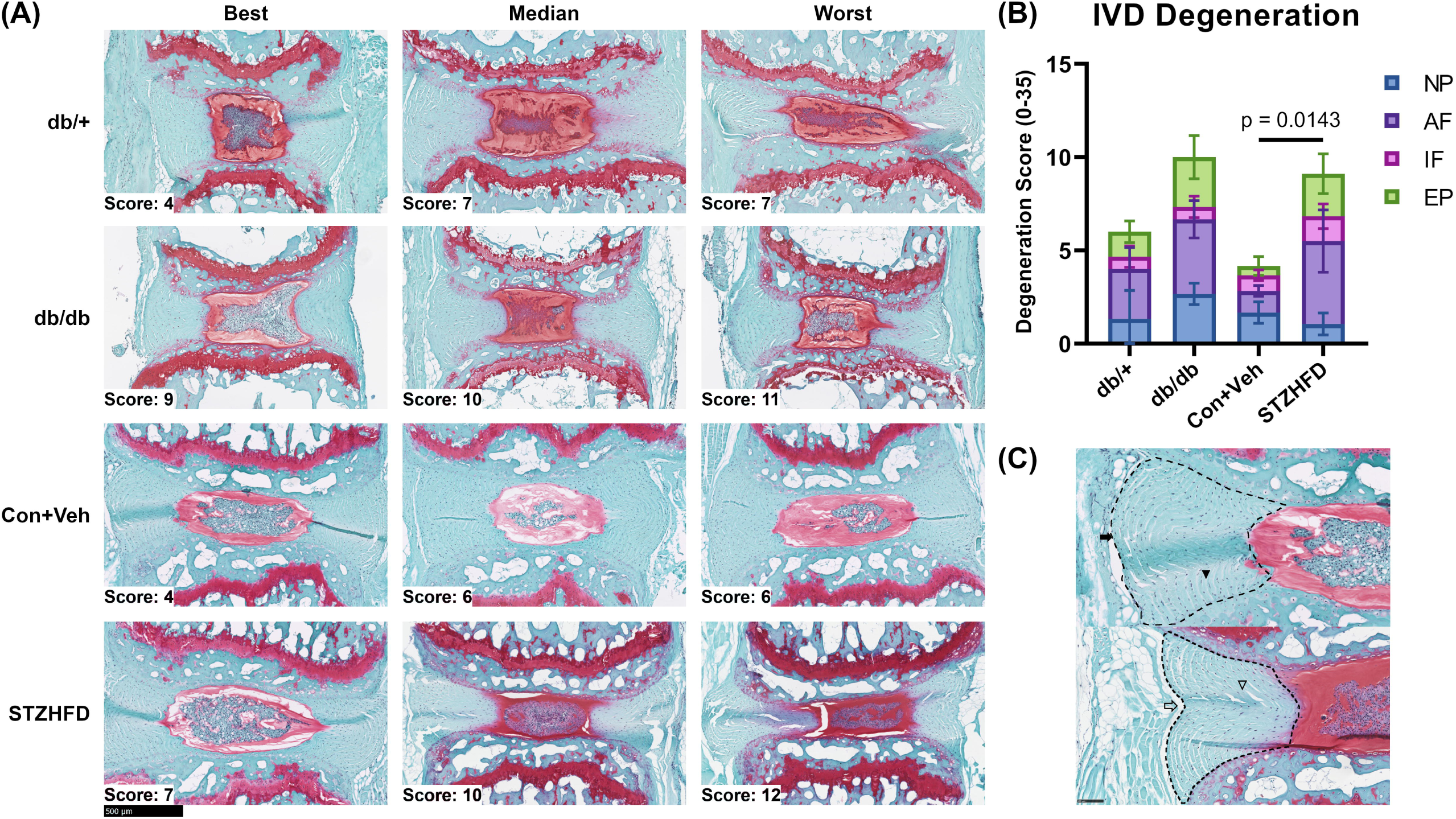
STZ-HFD mice exhibit more severe histopathological IVD degeneration compared to db/db mice. **(A)** Comparison of histological phenotypes in db/+, db/db, Con+Veh, and STZ-HFD groups, showing best, median, and worst samples. **(B)** Histopathological scoring shows significantly greater IVD degeneration in STZ-HFD mice versus controls. **(C)** Con+Veh samples have healthy lamellae (Ti+); STZ-HFD samples show degenerate, disorganized lamellae (¢V).

### CT Analysis, Matrix Assays, and Mechanical Testing All Show No Major Effects in T2D IVD

The analysis of CEµCT data, mechanical testing, and matrix protein assays revealed only one significant difference among the four groups across all nine measured outcomes. Specifically, parameters such as NI/DI, NPVF, loading slope, hysteresis energy, tan delta, and biochemical content showed no variations between groups (**Fig. 4 A-B, D-I**). The only statistically significant result was a difference in morphology between the db/db and STZ-HFD groups, as indicated by the DHI (**Fig. 4C**). These findings suggest that the structural integrity, mechanical behavior, and biochemical composition of the IVDs are mostly consistent across the db/db and STZ-HFD models.

**Figure 4.**
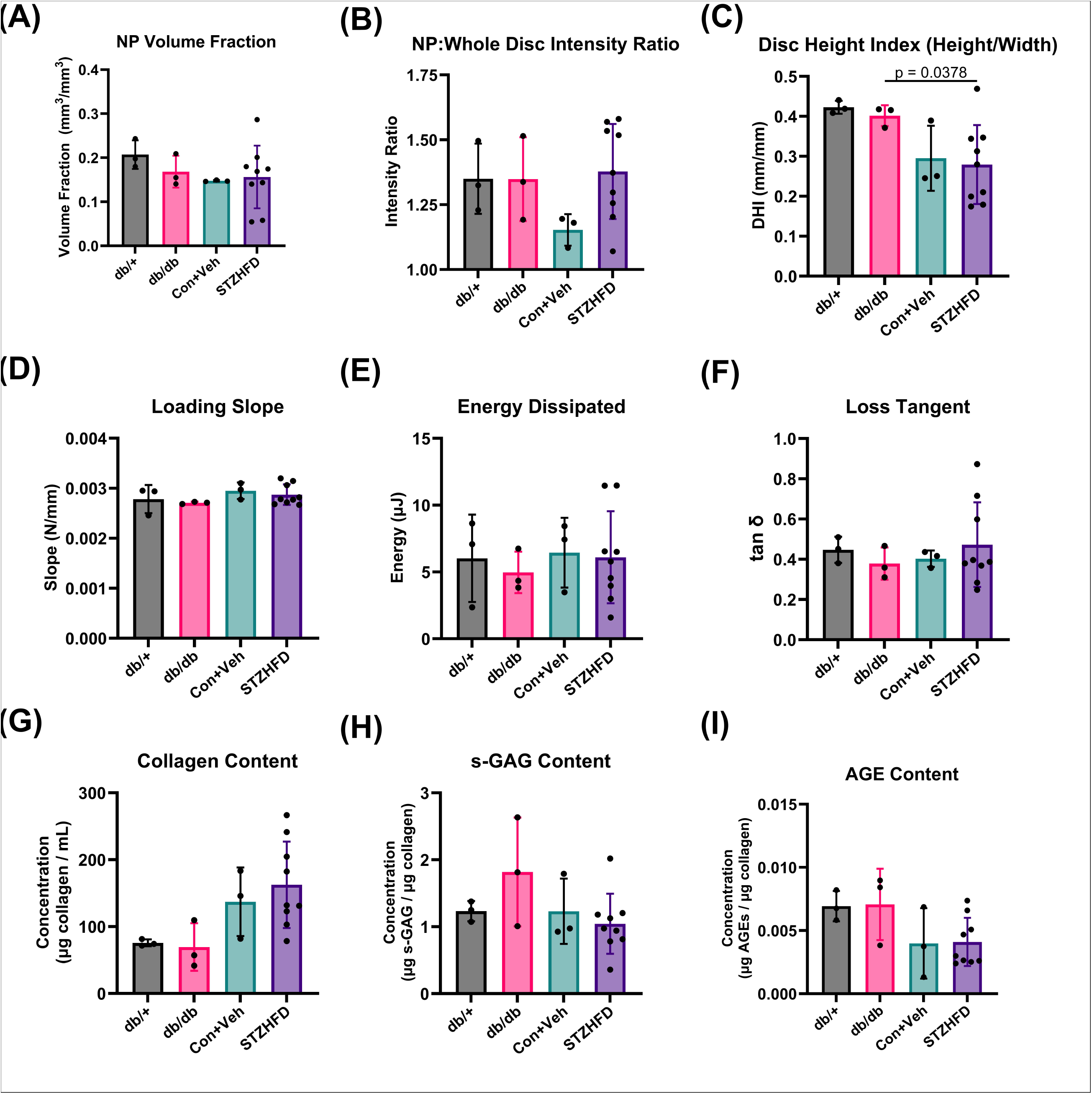
Comparative analysis of IVD structure, mechanics, and composition are similar between db/db and STZ-HFD mice. (A)-(C) NPVF, NIDI, and DHI indicate few significant differences in structural integrity between the models. **(D)-(F)** Load slope, energy dissipated, and phase shift demonstrating no significant variations in viscoelastic mechanical behavior. **(G)-(I)** Biochemical content measurements, including collagen, s-GAG, and AGEs, show no significant differences.

### Immunomodulatory Cytokines Are Chronically Up-regulated in STZ-HFD IVDs

The comparative analysis of cytokine expression levels between db/db and STZ-HFD models provides novel insights into diabetic inflammation of the IVD. The initial comparative analysis of the two models’ protein expression levels revealed that two cytokines (CCL2, CCL3) and sixteen cytokines (CCL2,3,4,5,12; CXCL1,2,9,10; CX3CL1; IL-2,6,16; CSF-3; VEGF; LIF) were up-regulated in the db/db and STZ-HFD models respectively, and thus included in the fold change analysis. The fold change in cytokine levels is computed over their respective controls (db/+ for db/db, Con+Veh for STZ-HFD) (**Fig. 5A**). The STZ-HFD model exhibited a significantly higher fold increase compared to the db/db model for 8 cytokines: CXCL2, CCL2, CCL3, CCL4, CCL12 (monocyte/macrophage associated cytokines) (Sagar et al., 2012; Arendt et al., 2013; Lança et al., 2013; Etna et al., 2014; Motwani and Gilroy, 2015; He et al., 2016; Lim et al., 2016; DeLeon-Pennell et al., 2017; Ruytinx et al., 2018; Sindhu et al., 2019; Zhang et al., 2019, 38; Huang et al., 2020; Pelisch et al., 2020; Yang et al., 2020; Xu et al., 2021a; Xu et al., 2021b; Sheng et al., 2022), IL-2, CXCL9 (T-cell associated cytokines) (Chang and Radbruch, 2007; Venetz et al., 2010; Shachar and Karin, 2013; Ochiai et al., 2015; Boff et al., 2018; Kuo et al., 2018; Mortara et al., 2018; House et al., 2020; Marcovecchio et al., 2021; Markovics et al., 2022), and CCL5 (pleiotropic cytokine) (Juhas et al., 2015; Atri et al., 2018; Kranjc et al., 2019; Chen et al., 2020; Zeng et al., 2022). Representing the overlapping cytokine expression profiles through a Venn diagram revealed that the STZ-HFD model encompassed a large number of up-regulated cytokines (**Figure 5B**). The inner circle represents the db/db model, containing only the two up-regulated cytokines, both of which are also up-regulated in the STZ-HFD model. This indicates that the STZ-HFD model has a broader and more pronounced cytokine response compared to the db/db model.

**Figure 5.**
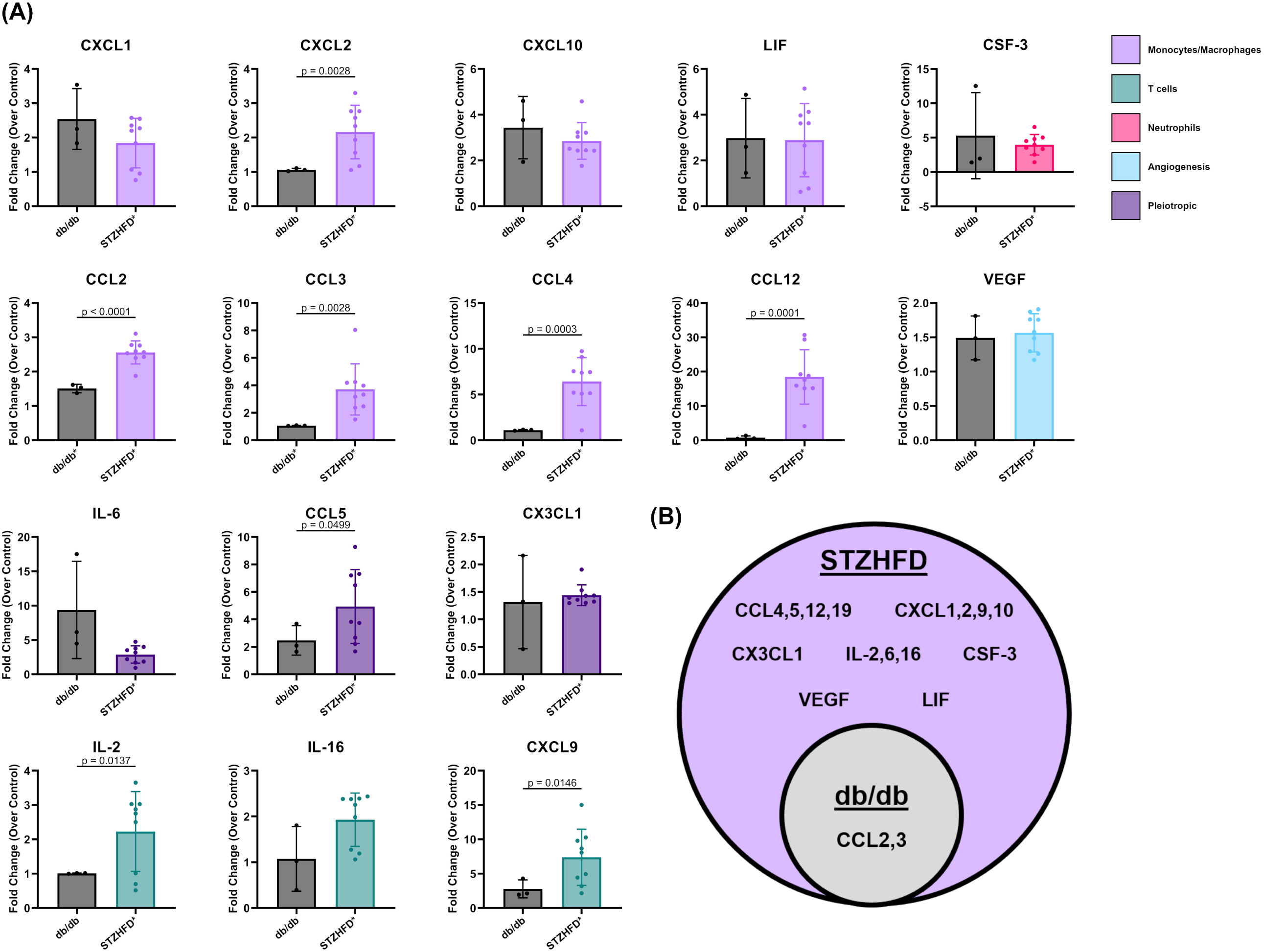
STZ-HFD IVD Produces a More Pro-Inflammatory Microenvironment than the db/db IVD in Comparative Analysis of Cytokine Expression. **(A)** The STZ-HFD model shows a significantly higher fold increase for eight cytokines: CXCL2, CCL2, CCL3, CCL4, CCL12 (monocyte/macrophage associated), IL-2, CXCL9 (T-cell associated), and CCL5 (pleiotropic). **(B)** The STZ-HFD model encompasses a broader and more pronounced cytokine response compared to the db/db model, highlighting the extensive up-regulation of inflammatory cytokines in the STZ-HFD model.

### Differential Network Structures Reveal Unique Inflammatory Pathways of T2D in IVD

Our cytokine network analysis provided detailed insight into how inflammation is organized and coordinated within the IVD microenvironment of each T2D model (**Fig. 6A**). Key cytokines such as CCL2 and CCL4 appeared centrally connected in both models, acting as critical communication hubs that likely coordinate broader inflammatory responses, as indicated by high betweenness centrality (**Fig. 6B**). When examining each model individually, the db/db network featured several unique cytokines (CSF3, CXCL5, CXCL9, CXCL10, IL-4, and IL-11) that held prominent positions in terms of connectivity and influence (betweenness and eigenvector centralities). These cytokines may reflect pathways uniquely modulated by leptin deficiency. In contrast, the STZ-HFD model showed a distinct set of highly connected and influential cytokines, including CXCL2, IL-6, IL-16, CCL11, and CSF3, suggesting alternative signaling axes are activated when leptin signaling remains intact. The organization of these networks also differed in terms of their communication efficiency: the db/db model exhibited a shorter average path length, meaning inflammatory signals can propagate more quickly and uniformly. The STZ-HFD network had longer path lengths and higher modularity, indicating a more compartmentalized structure with distinct inflammatory sub-networks, likely reflecting a more complex and widespread inflammatory state (Fig. 6C). To evaluate the extent of overlap between models, we used the Jaccard index to assess network similarity. The low similarity between STZ-HFD and WT controls (Jaccard index: 0.144 at k=1, 0.246 at k=2) confirms a substantial remodeling of cytokine interactions in this model. Conversely, the db/db model’s network remained more similar to its control, indicating a less dramatic inflammatory shift.

**Figure 6.**
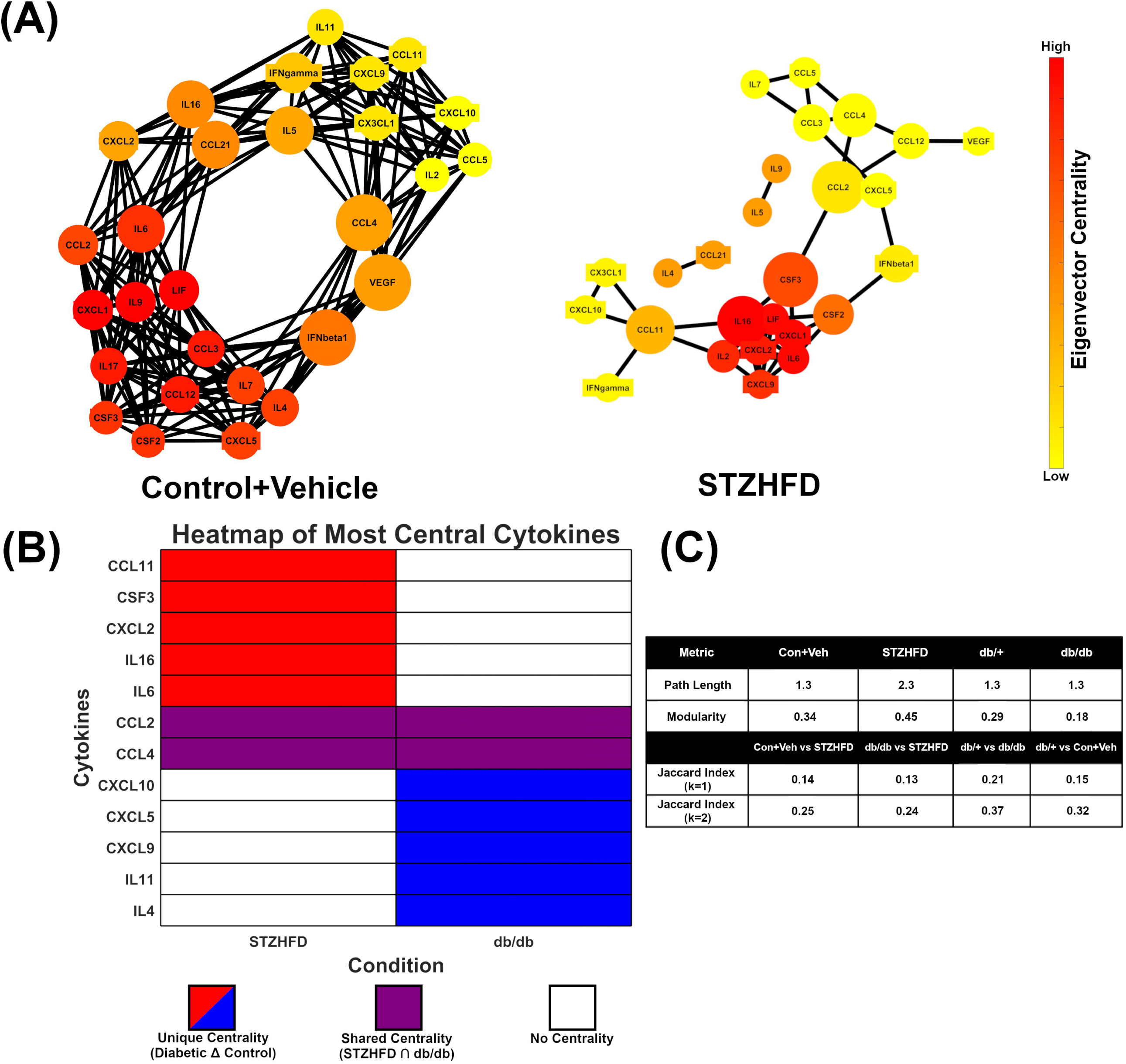
STZ-HFD IVD Invokes Unique Inflammatory Signaling Pathways in Networks of Cytokine Expression. **(A)** The STZ-HFD model shows a distinct network structure, demonstrating the unique up-regulation of various inflammatory pathways **(B)** The STZ-HFD and db/db networks each rely on a number of unique (red/blue) and shared (purple) cytokines, indicating both leptin-dependent and leptin-independent inflammatory signaling cascades **(C)** The STZ-HFD mouse model displays a fragmented and modular cytokine network, indicating the parallel signaling of multiple signaling pathways.

Together, these findings suggest that although both models exhibit inflammation, the STZ-HFD model engages broader and more structured cytokine signaling cascades, better representing the diffuse and chronic nature of T2D-associated disc degeneration.

### STZ-HFD IVDs Show Distinct Gene Expression and Pathway Dysregulation in RNA-Seq Analysis

The RNA-seq analysis revealed significant differences between the db/db and STZ-HFD models in several biological processes, particularly those related to type 2 diabetes-related inflammatory pathways, immune response, ECM organization, metabolic function, and signal transduction. Both models had many genes up-regulated and down-regulated when adjusting for false discovery rate and limiting to genes with high absolute fold change (**Fig. 7A**). The db/db model exhibited 376 up-regulated genes, while the STZ-HFD model exhibited 170; 3 genes (*Adamts8, Cd36, Ndufb5*) were shared between the two models (**Fig. 7B**). Likewise, the db/db model showed 85 down-regulated genes, while the STZ-HFD model showed 135, with 1 gene (*Glis1)* shared between the models. Eighteen genes associated with the metabolic processes of glycolysis, sugar reduction, and carbonyl metabolism were differentially expressed between the db/db and STZ-HFD models (**Fig. 7C**); these intracellular mechanisms are critical in Type 2 Diabetes pathology through the formation of AGEs and activation of the receptor for AGEs (RAGE). In the db/db model, 5 genes were up-regulated and 4 were down-regulated, whereas in the STZ-HFD model, 5 genes were up-regulated and 7 were down-regulated. Additionally, several damage-associated molecular patterns (DAMPs) and RAGE-modulating genes were changed (**Fig. 7C**). Specifically, 4 genes were up-regulated in the db/db model, while 6 genes were down-regulated in the STZ-HFD model. Furthermore, regulators of NF-κB signaling, a pathway implicated in inflammatory responses downstream of RAGE signaling, was moderately affected. In the db/db model, 1 gene was up-regulated and 1 down-regulated, while in the STZ-HFD model 1 gene was up-regulated and 3 genes were down-regulated (**Fig. 7C**). Regarding adipokine signaling, which is critical in the regulation of glucose and lipid metabolism in Type 2 Diabetes, 2 genes related to leptin signaling were differentially expressed in the STZ-HFD model while only 1 gene was differentially expressed in the db/db model (**Fig. 7C**). Additionally, the db/db model demonstrated an increase in the adiponectin-related gene *Adipoq*. This ultimately strengthens our initial hypothesis regarding the role of leptin signaling in mediating diabetic inflammation and degeneration of the IVD.

**Figure 7.**
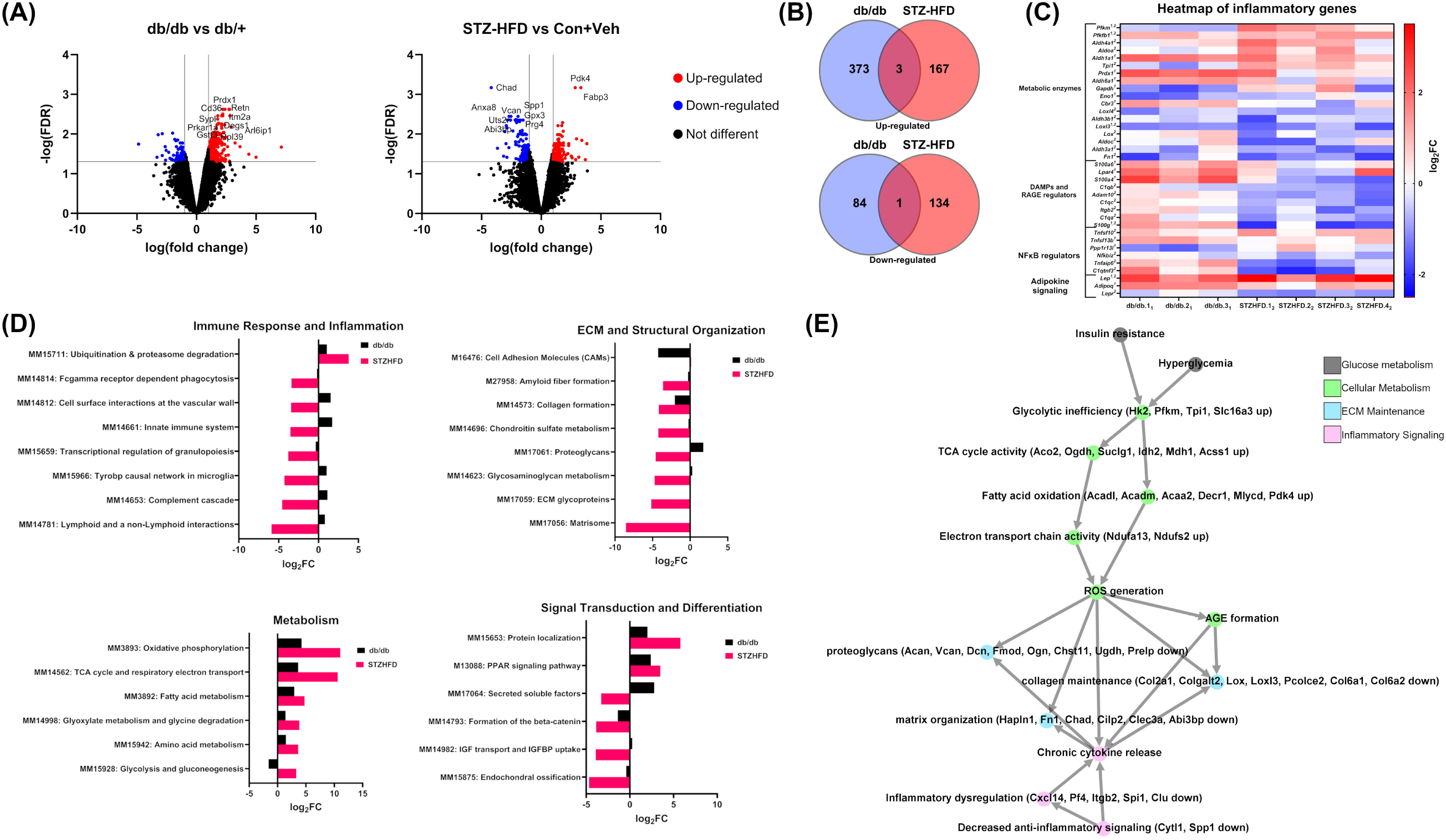
Enhanced Pathway Disruption in the STZHFD Model Highlights the Complexity of T2D Pathophysiology in the IVD. **(A)** A volcano plot displaying the significantly up-regulated genes. Several of the most up-regulated genes to appear in later ontological analyses have been labeled for the STZ-HFD model. **(B)** Venn diagrams comparing up- and down-regulated genes in the db/db model and the STZ-HFD model. Only 3 up-regulated genes and 1-down regulated gene was shared between the two models. **(C)** The STZ-HFD (marked with 2) model exhibits greater transcriptional alterations in T2D-related pathways than db/db (marked with 1) model, particularly in non-enzymatic glycosylation and RAGE signaling. **(D)** Immune response and inflammation pathways, including ubiquitination and proteasome degradation, are significantly disrupted in the STZ-HFD model but remain unaltered in db/db. ECM remodeling and structural integrity pathways are more extensively down-regulated in the STZ-HFD model, suggesting greater tissue disruption. Metabolic dysfunction is more pronounced in the STZ-HFD model, with additional up-regulation of pathways involved in amino acid metabolism and gluconeogenesis. Signal transduction and tissue differentiation pathways show significant alterations in the STZ-HFD model, indicating complex changes in cellular communication and development processes. **(E)** Based on ontological pathway analysis and significantly up-regulated genes, a suggested mechanistic diagram of T2D-induced intervertebral disc degeneration in the STZ-HFD mouse has been provided. Groups of processes have been grouped as diabetic symptoms, metabolic changes, immune/inflammatory changes, and matrix changes.

Enrichment analysis identified eight pathways in the Molecular Signatures Database (MSigDB, https://www.gsea-msigdb.org/gsea/msigdb) significantly affected in the STZ-HFD model, all of which are crucial for immune response and inflammation. These included the up-regulation of ubiquitination and proteasome degradation, and the down-regulation of pathways such as Fcgamma receptor-dependent phagocytosis, cell surface interactions at the vascular wall, innate immune system, transcriptional regulation of granulopoiesis, the Tyrobp causal network in microglia, complement cascade, and lymphoid/non-lymphoid interactions (**Fig. 7D**). Interestingly, none of these pathways were significantly altered in the db/db model, highlighting a distinct inflammatory and immune regulation in the STZ-HFD model.

Eight MSigDB pathways related to ECM and structural organization were significantly altered between the models. In the db/db model, the pathway associated with cell adhesion molecules was down-regulated. Conversely, in the STZ-HFD model, several pathways were down-regulated, including amyloid fiber formation, collagen formation, chondroitin sulfate metabolism, proteoglycans, glycosaminoglycan metabolism, ECM glycoproteins, and the matrisome (**Fig. 7D**). This suggests that ECM remodeling and structural integrity are more disrupted under STZ-HFD conditions compared to db/db at the transcriptional level.

The analysis also revealed six MSigDB pathways significantly impacted by metabolic function. Both models showed up-regulation in oxidative phosphorylation, TCA cycle and respiratory electron transport, and fatty acid metabolism. However, additional pathways were specifically up-regulated in the STZ-HFD model, including glyoxylate metabolism and glycine degradation, amino acid metabolism, and glycolysis and gluconeogenesis (**Fig. 7D**). This indicates a greater depth to the metabolic dysfunction in the STZ-HFD model relative to db/db.

Finally, six MSigDB pathways were identified as being significantly involved in signal transduction and tissue differentiation. In the STZ-HFD model, pathways related to protein localization and PPAR signaling were up-regulated, while pathways involved in secreted factors, the formation of beta-catenin complexes, IGF transport and IGFBP uptake, and endochondral ossification were down-regulated (**Fig. 7D**). These findings suggest alterations in signaling mechanisms and tissue differentiation processes that are more pronounced in the STZ-HFD model compared to db/db. Overall, the STZ-HFD model exhibited more extensive and varied disruptions across multiple canonical biological pathways, as well as type 2 diabetes-related genes and processes, including those involved in glycosylation, immune response, ECM structure, metabolism, and signal transduction, indicating its utility in studying complex metabolic and inflammatory disorders compared to the db/db model.

## Discussion

This study offers detailed comparative analyses of chronic inflammatory profiles and IVD degeneration in murine T2D models, specifically contrasting the db/db and STZ-HFD mouse models. Our key findings indicate that while both models exhibit hallmark systemic diabetic phenotypes (Cefalu, 2006; Dalgaard and Pedersen, 2001; Lin and Sun, 2010), the STZ-HFD model demonstrates more pronounced IVD degeneration, greater matrix disruption, and broader transcriptional dysregulation of inflammatory pathways. This suggests that the STZ-HFD model as a more physiologically relevant model for studying T2D-associated musculoskeletal complications due to inflammation (Gilbert et al., 2011; Stott and Marino, 2020). By integrating mechanical testing, histology, transcriptomics, and cytokine profiling, our study directly addresses the open questions identified in the introduction regarding how T2D alters inflammatory signaling, disc structure, and mechanical function. These findings support our hypothesis that the STZ-HFD model better recapitulates the inflammatory and degenerative features of T2D within the IVD, providing a foundational platform for future mechanistic investigations and therapeutic testing.

In this study, we induced a consistent T2D phenotype in C57BL/6 mice using HFD and a single dose of 100 mg/kg STZ, causing pronounced chronic hyperglycemia, insulin resistance, and obesity. This phenotype has been extensively characterized in prior studies, including Eckhardt et al. (2020), who used the identical diet and STZ regimen and reported hallmark T2D features such as elevated fasting blood glucose, increased A1C, and substantial adiposity without the significant β-cell loss characteristic of T1D. Similarly, Gao et al., 2024, demonstrated that this regimen induces hyperglycemia, insulin resistance, hyperinsulinemia, and elevated A1C, aligning with a metabolic profile consistent with T2D. Additional studies (Barrière et al., 2018; Luo et al., 1998; Mazzocco et al., 2025; Racine et al., 2024; Yin et al., 2020) using comparable STZ-HFD protocols have confirmed similar outcomes, underscoring the reproducibility and translational potential of this model. In our own validation, we performed an insulin tolerance test (Supplemental Figure S2) showing that STZ-HFD animals exhibited poor glucose clearance in response to exogenous insulin, a key feature of insulin resistance. Notably, this profile differs from T1D models, where insulin deficiency leads to dramatic glucose reductions following insulin administration. Taken together, the metabolic profile observed in our study strongly supports the conclusion that the STZ-HFD model used here recapitulates the physiological characteristics of T2D and is therefore an appropriate model for investigating T2D-related intervertebral disc pathology.

In terms of IVD degeneration, our study demonstrates that histopathologically STZ-HFD mice exhibit significantly greater IVD degeneration compared to controls, indicating a notable degenerative phenotype. Assessment of structural integrity, mechanical behavior, and biochemical composition did not reveal further significant differences, however. In contrast, db/db mice, despite severe obesity, insulin resistance, and chronic hyperglycemia, show milder IVD degeneration compared to littermate controls. This observation aligns with existing literature which reports moderate IVD degeneration in db/db mice influenced by variables such as sex and specific metabolic disruptions (Li et al., 2020; Lintz et al., 2022; Natelson et al., 2020).

To distinguish the inflammatory mechanisms relevant to T2D-induced IVD degeneration, our findings demonstrate that the STZ-HFD model exhibits a broader and more diversified cytokine profile within the IVD compared to the db/db model. Specifically, STZ-HFD mice showed significant up-regulation of 16 cytokines, including those involved in monocyte and macrophage recruitment (e.g., CCL2, CCL3, CXCL2), T-cell activation (e.g., IL-2, CXCL9), and broader immune modulation (e.g., CCL4, CCL5, IL-6, IL-16, CSF3). These cytokines mirror inflammatory mediators frequently elevated in human T2D patients, supporting this model’s utility in investigating immune-mediated tissue degeneration linked to metabolic dysfunction (Alshammary et al., 2023; Chang et al., 2021; Inayat et al., 2019; Keophiphath et al., 2010; Mir et al., 2024, 2; Pan et al., 2021; Pettigrew et al., 2010). In contrast, the db/db model produced a more limited cytokine expression profile, shaped in part by the absence of leptin signaling, which modulates both innate and adaptive immune responses (Francisco et al., 2018). While the db/db model remains informative for studying hyperglycemia and leptin-deficiency-related disc changes, its restricted inflammatory activation underscores its limitations in modeling the broad cytokine dysregulation observed in most human T2D cases. By comparing these models, we were able to address specific questions regarding the contribution of intact leptin signaling to IVD inflammation and identify cytokines that may play mechanistic roles in disc degeneration under conditions of metabolic dysfunction.

Cytokine network analysis revealed distinct patterns of cytokine interactions in the STZ-HFD IVD, highlighting differences in regulatory dynamics and pathway activation compared to the db/db model. The STZ-HFD network demonstrated greater modularity and longer average path length, indicating compartmentalized signaling across multiple, distinct inflammatory circuits. This potentially reflects the multifactorial nature of T2D pathophysiology, which includes contributions from obesity, insulin resistance, and metabolic inflammation (Francisco et al., 2018; Pan et al., 2021; Rebuffat et al., 2018; Segar et al., 2019; Sharma, 2018). Central cytokines in this model included CXCL2, IL-6, IL-16, CCL11, and CSF3, several of which are known mediators of monocyte recruitment and chronic inflammation in diabetic complications, suggesting their possible involvement in IVD degeneration and potential as therapeutic targets. In comparison, the db/db model exhibited a distinct inflammatory signature, with central cytokines such as CSF3, CXCL5, CXCL9, CXCL10, IL-4, and IL-11. This divergence may reflect the impact of leptin receptor deficiency on immune regulation, as leptin is known to influence both pro- and anti-inflammatory cytokine expression and immune cell activation (Francisco et al., 2018). These findings suggest that while both models exhibit cytokine-driven inflammation, the signaling architecture differs substantially, with db/db mice exhibiting a profile consistent with impaired leptin-mediated immune modulation.

The RNA-seq data reveals significant transcriptional alterations in pathways related to RAGE-driven inflammation within the IVD in the STZ-HFD model. This model showed an increase in phosphofructokinases (*Pfkm, Pfkfb1*) and a triosephosphate isomerase (*Tpi1*), which ultimately promote the accumulation of glycolytic intermediates that readily form AGE precursors such as methylglyoxal (Gonzalez et al., 1980; Hamada et al., 1996; Hellman, 1970; Twarda-Clapa et al., 2022), that may subsequently enrich AGE formation long-term.

Additionally, changes in DAMP production and regulators of RAGE signaling indicate a shift in favoring sugar-derived AGEs in the STZ-HFD model. Specifically, the STZ-HFD IVD heavily down-regulates several DAMPs, which are alternative ligands for RAGE, likely in an attempt to reduce binding competition with the AGEs (Chavakis et al., 2003; Hofmann et al., 1999). In parallel with this, the down-regulation of *Itgb2* dampens non-AGEs-RAGE binding and the suppression of *Adam10* increases retention of membrane-bound RAGE (Chavakis et al., 2003; Lee et al., 2015). Directly down-stream of RAGE is the broad inflammatory transcription factor NF-κB. In the STZ-HFD IVD, elevation of *Tnfsf6* and decrease in *Tnfaip6* point towards an increase in NF-κB activation, in alignment with our speculation that RAGE and its downstream targets play a role in promoting inflammation (Hayden and Ghosh, 2014; Lai et al., 2013). These findings suggest molecular reprogramming within the diabetic IVD that favors AGE-RAGE signaling over DAMP-mediated pathways, possibly tipping the balance toward sustained NF-κB activation and inflammation. Our analysis of secreted chemokines corroborates this finding as many of the chemokines are downstream of NF-κB.

As observed in literature, genetic deletion of RAGE in murine models significantly attenuates intervertebral disc degeneration by preserving collagen structure, reducing AGE accumulation, and lowering expression of pro-inflammatory mediators and matrix-degrading enzymes, indicating that RAGE signaling directly mediates AGE-induced structural and inflammatory damage within the disc (Hoy et al., 2020; Walk, 2024). Similarly, inhibition of NF-κB activity in rodent IVDs protects against age-related and mechanically induced disc degeneration by reducing cytokine and MMP expression, maintaining extracellular matrix integrity, and improving histological disc scores (Burt et al., 2024; Glaeser et al., 2020; Li et al., 2023; Nasto et al., 2012; Orita et al., 2013).

Canonical pathway analysis further emphasizes the distinct biological functions altered in these T2D models. In the STZHFD model, the significant up-regulation of ubiquitination and proteasome degradation pathways, coupled with the down-regulation of immune-related pathways within the IVD, indicate a stress response aimed at managing protein damage, reflecting the chronic inflammatory state associated with metabolic diseases like T2D (Garcia-Martinez et al., 2015). The down-regulation of ECM and structural organization pathways, including collagen formation and ECM glycoproteins, hints at impairment of tissue remodeling and fibrosis within the IVD, aligning with the structural changes observed in the histopathological analysis of the STZHFD IVD. In contrast, metabolic pathways such as oxidative phosphorylation and fatty acid metabolism were up-regulated in both models, reflecting increased metabolic demands. However, the STZHFD model showed additional up-regulation in pathways like glyoxylate metabolism and glycolysis/gluconeogenesis, indicating more extensive metabolic reprogramming within the IVD, further illustrating the model’s ability to capture the systemic nature of T2D-related complications (Joshi et al., 2020; Dhawan et al., 2022).

The gene ontology analysis identified the most significantly changed genes in these pathways, and constructed a mechanistic diagram in an attempt to understand the potential molecular relationships behind T2D-induced IVD degeneration in the STZ-HFD model (**Fig. 7E**). It is likely that the combination of insulin resistance and hyperglycemia leads to abnormal intracellular glucose concentrations, which in turn leads to glycolytic inefficiency. This is evidenced by the up-regulation of *Hk2*, *Pfkm*, *Tpi1*, and *Slc16a3* (all related to progressing glycolysis). This inefficiency has two parallel effects in IVD cells. Firstly, TCA cycle activity, marked by increases in *Aco2*, *Ogdh*, *Suclg1*, *Idh2*, *Mdh1*, and *Acss1,* compensate for disrupted cellular energetics. Downstream, this produces more NADH in the mitochondria, which pass through the electron transport chain down the proton gradient, marked by increases in *Ndufa13* and *Ndufs2*. Secondly, in parallel to these processes, fatty acids are shunted into β-oxidation, evidenced by increases in *Acadl*, *Acadm*, *Acaa2*, *Decr1*, *Mlycd*, and *Pdk4*. Both of these metabolic processes result in excess reactive oxygen species (ROS) being generated, enhancing the kinetics of AGEs formation. In parallel to metabolic changes, we observed a decrease in anti-inflammatory genes (*Cytl1* and *Spp1*) and cytokine regulators (*Cxcl14*, *Pf4*, *Itgb2*, *Spi1*, and *Clu*). This is corroborated by the enhanced cytokine release observed in the secretome of the STZ-HFD IVDs. Changes in both the metabolic and inflammatory regimes in these cells converge to disrupt the vital processes for maintaining a healthy IVD matrix. Primarily, there were decreases in genes related to proteoglycan production (*Acan*, *Vcan*, *Dcn*, *Fmod*, *Ogn*, *Chst11*, *Ugdh*, *Prelp*), collagen maintenance (*Col2a1*, *Colgalt2*, *Lox*, *Loxl3*, *Pcolce2*, *Col6a1*, *Col6a2*), and matrix organization (*Hapln1*, *Fn1*, *Chad*, *Cilp2*, *Clec3a*, *Abi3bp*).

Despite these significant findings, this study has several limitations. First and foremost, this study exclusively evaluates diabetes in male mice due to limitations of the STZ-HFD model. While many symptoms and mechanisms of diabetes are shared between males and females (in mice as well as in humans), there are important sex-specific effects to be studied still. Namely, women with diabetes experience disproportionately higher risks of complications such as cerebral microvascular disease leading to cognitive decline, greater incidence of coronary heart disease, and heightened vascular dysfunction, particularly post-menopause (Kautzky-Willer et al., 2023; Peters et al., 2014; Thomas et al., 2022). These distinctions suggest that hormonal regulation and immune-metabolic interactions in females may critically influence diabetes pathology and should be explored in future studies on female mice by using a higher dose of STZ or ovariectomized mice to more consistently induce diabetes. Secondarily, the selection of cytokines examined was relatively small, potentially missing other important inflammatory mediators involved in IVD degeneration. Additionally, the mechanistic link between inflammation and degeneration remains unclear and warrants further investigation. While the STZ-HFD model provides a comprehensive inflammatory profile, the specific pathways driving the observed IVD degeneration need to be elucidated through future studies to appropriately identify therapeutic targets. Finally, the db/+, db/db, and Con+Veh groups in this study may be underpowered due to small sample size, and thus this study most thoroughly characterizes the STZ-HFD model in the metabolic and disc-specific assays. Future directions for research based on our findings include expanding the panel of cytokines and other inflammatory mediators examined in the STZ-HFD model to gain a more complete understanding of the inflammatory landscape in T2D. Investigating the specific molecular and cellular mechanisms linking inflammation to IVD degeneration will be crucial to furthering future therapeutic approaches. Finally, exploring therapeutic interventions targeting the identified cytokine pathways could provide insights into potential treatments for T2D-related IVD degeneration. While the STZHFD model showed down-regulation of ECM-related pathways at the mRNA level, compensatory translational mechanisms may prevent a corresponding reduction in ECM proteins, and the modest changes in NF-κB signaling underscore the limitations of transcriptomic data alone (Mobeen and Ramachandran, 2020; Wang et al., 2020). Consequently, integrating protein measurements with transcriptomic analyses is essential for fully understanding the molecular processes of the T2D IVD.

Our findings establish the STZ-HFD model recapitulates the clinical definitions for T2D diabetes, and the inflammatory profiles of these IVDs may be suitable for T2D-related mechanisms in IVD degeneration. The extensive cytokine up-regulation and significant degenerative phenotype observed in this model provide a framework for T2D-associated pathologies. The db/db model, while still relevant, exhibits a more stunted cytokine profile and limited IVD degeneration. These insights enhance our understanding of T2D-induced IVD degeneration and identify key cytokine pathways for therapeutic development, emphasizing the need to address these pathways holistically for effective intervention.

## Acknowledgements

Thank you to the Alafi Neuroimaging Lab for Nanozoomer Access (NIH S10RR027552).

Multiplex cytokine panels were performed by Eve Technologies Corp.

---

**Figure S1.** Representative force-displacement curve of the dynamic loading regimen used in mechanical testing. The force-displacement curve here illustrates the loading hysteresis in these samples, consistent with behavior of a viscoelastic material.

**Figure S2.** Insulin tolerance test of STZ-HFD mice shows insulin resistance, not insulin dependence. Despite receiving 2U/kg I.P. insulin (Humalin R), STZ-HFD mice do not respond with a reduction in blood glucose concentration. Consequently, STZ-HFD mice have elevated blood glucose when compared to non-diabetic controls at all timepoints during the insulin tolerance test.

