## Supplementary figures and images for "The transcriptome and secreted factors of the intervertebral discs in STZ-HFD Type□2 diabetic male mice reveal extensive inflammation"

### Supplement Figure S1

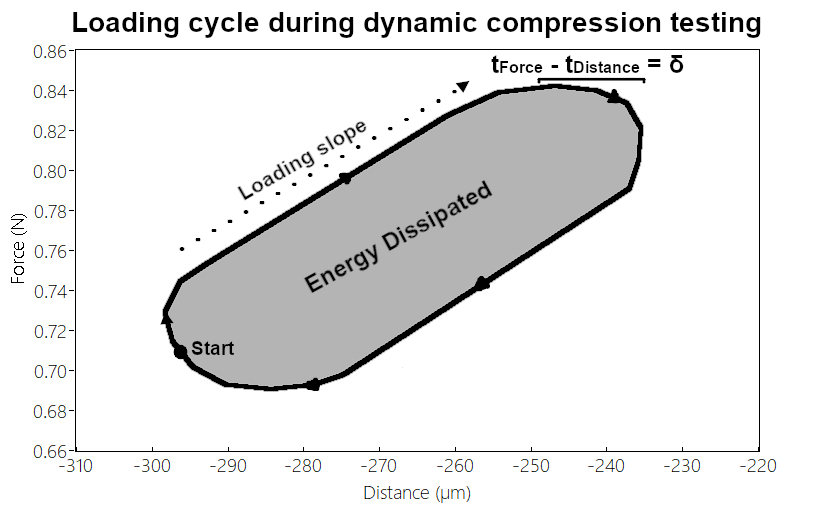

### Supplement Figure S2

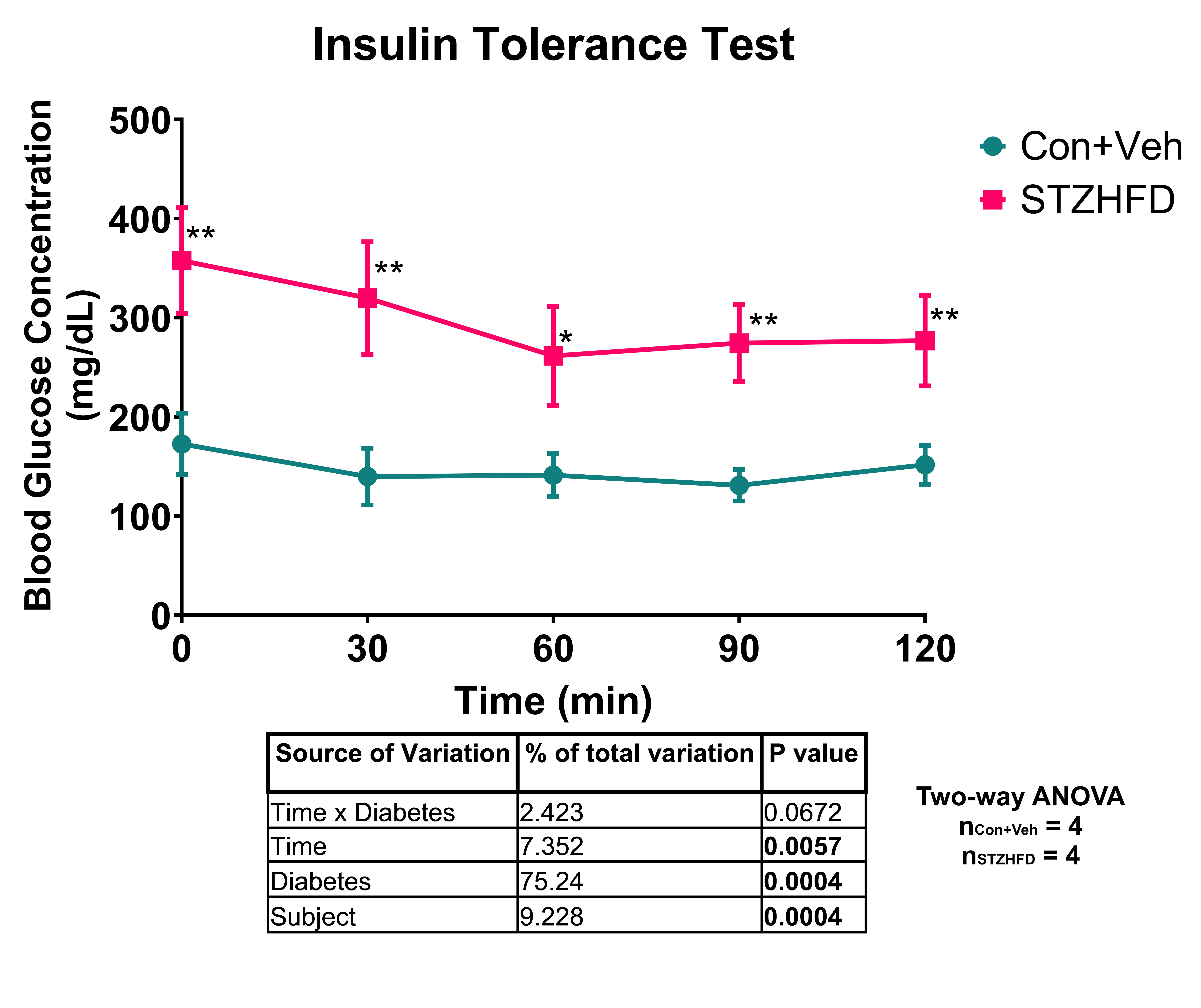
